# Power-law scaling of brain wave activity associated with mental fatigue

**DOI:** 10.1101/2020.08.03.234120

**Authors:** Vo V. Anh, Hung T. Nguyen, Ashley Craig, Yvonne Tran, Yu Guang Wang

## Abstract

This paper investigates the cause and detection of power-law scaling of brain wave activity due to the heterogeneity of the brain cortex, considered as a complex system, and the initial condition such as the alert or fatigue state of the brain. Our starting point is the construction of a mathematical model of global brain wave activity based on EEG measurements on the cortical surface. The model takes the form of a stochastic delay-differential equation (SDDE). Its fractional diffusion operator and delay operator capture the responses due to the heterogeneous medium and the initial condition. The analytical solution of the model is obtained in the form of a Karhunen-Loève expansion. A method to estimate the key parameters of the model and the corresponding numerical schemes are given. Real EEG data on driver fatigue at 32 channels measured on 50 participants are used to estimate these parameters. Interpretation of the results is given by comparing and contrasting the alert and fatigue states of the brain.

The EEG time series at each electrode on the scalp display power-law scaling, as indicated by their spectral slopes in the low-frequency range. The diffusion of the EEG random field is non-Gaussian, reflecting the heterogeneity of the brain cortex. This non-Gaussianity is more pronounced for the alert state than the fatigue state. The response of the system to the initial condition is also more significant for the alert state than the fatigue state. These results demonstrate the usefulness of global SDDE modelling complementing the time series approach for EEG analysis.

## 1. Introduction

Mental fatigue is a significant cause of accidents and injury in driving [11] and in performing repetitive tasks, in-process works [1]. There have been many studies undertaken to determine the association between mental fatigue and brain activity. These studies mostly employed electroencephalography (EEG) to measure brain activity and examined the changes in the EEG as a person moves from the alert state to a fatigue state. Table 1 of [12] presents a summary of 17 such studies and their findings. The changes in the EEG were commonly detected by computing the fast Fourier transform of the EEG time series and analysing transformed data at the following frequency bands: delta wave (0.5 to 3.5 Hz), theta wave (4 to 7.5 Hz), alpha wave (8 to 13 Hz) and beta wave (14 to 30 Hz). Some main findings from the above studies include that there seem to be no significant delta wave changes associated with fatigue, theta and alpha wave activities increase significantly during fatigue, but where, in the cortex, these changes occur is still to be probed, and the association between beta wave activity and fatigue remains unclear. [29] probed into the delta and beta frequency bands to verify the existence of power-law scaling, usually realised in the form of 1*/f* -power spectrum. The authors of [29] used irregularly resampled auto-spectral analysis in conjunction with ARMA modelling to quantify the 1*/f* -component of magnetoencephalography, electroencephalography and electrocorticography (MEG/EEG/ECoG) power spectra in the low (0.1 to 2.5 Hz) and high (5 to 100 Hz) frequency bands. Their findings confirm power-law scaling in the MEG/EEG/ECoG in a more refined form of 1*/f* ^*β*^-power spectrum. Furthermore, the results follow a spatial pattern in the sense that, in the higher frequencies, steeper slopes are present in posterior areas. In contrast, for the lower frequencies, steeper slopes are present in the frontal cortex.

**Table 1:** Averages and standard deviations of the parameters *α, α* + *γ, τ* and *θ* over up to 50 participants

| Average | $\alpha$ | $\alpha + \gamma$ | $\tau$ | $\theta$ |
| --- | --- | --- | --- | --- |
| Alert | $0.921 \pm 0.207$ | $1.676 \pm 0.141$ | $0.183 \pm 0.061$ | $2.323 \pm 0.042$ |
| Fatigue | $1.116 \pm 0.208$ | $1.773 \pm 0.132$ | $0.160 \pm 0.048$ | $2.334 \pm 0.041$ |

Many complex systems in nature, from earthquakes to avalanches, are characterised by scale invariance, which is usually identified by a power-law distribution of variables such as event duration or the waiting time between events [6, 23]. The 1*/f* -noise is considered to be a footprint of such systems. 1*/f* -frequency scaling is the behaviour of a system near critical points. As such, one commonly associates self-organised critical states of a natural system with 1*/f* -frequency scaling [23]. Apart from MEG/EEG/ECoG, temporal signals displaying power-law scaling have been observed in many works on the nervous system at various spatial scales, from membrane potentials [16] to functional magnetic resonance imaging [20]. Despite its potential importance, the physiological mechanism which generates power-law scaling is still not well understood, and its significance for brain activity remains controversial [10]. It has been argued [9] that the existence of power-law scaling indicates that the brain is in a state of self-organised criticality. [8] pointed out that, alternatively, 1*/f* -frequency scaling may be due to the diffusion of EEG signals through various extracellular media such as cerebrospinal fluid, dura matter, cranium muscle and skin. Such a heterogeneous medium induces a combination of resistive effects, due to the flow of current in a conductive fluid, and capacitive effects due to the high density of membranes. In [7], the authors showed theoretically that 1*/f* -power spectra could be created by ionic current flow in such a complex network of resistors and capacitors with random values.

This paper will contribute a new angle to the debate on the cause and detection of power-law scaling of brain wave activity. We focus on the quantification of the cause, and subsequent response of the system, which models and interprets the heterogeneity of the brain cortex. Our starting point is the construction of a mathematical model of global brain wave activity based on all EEG measurements. Instead of modelling wave activities at various locations or regions of the brain, we will consider the evolution of the random field representing EEG over the entire cortical surface. The model takes the form of a stochastic delay-differential equation (SDDE) for this random field. Its two main components, the fractional diffusion operator and the delay operator capture respectively the two critical features of the system: the response due to the heterogeneous medium and the response to an initial condition, such as the alert or fatigue state of the brain. We will show that these two components are capable of generating respectively asymptotic temporal correlation and oscillatory rhythms of global EEG. The exponents of the fractional diffusion operator depict the effect of the heterogeneous medium on the diffusion. Thus, we build the power-law scaling which is due to heterogeneity of the medium into diffusion, as predicted by the theory of [7]. Another advantage of the model is that the asymptotic temporal correlation indicating power-law scaling is captured by the same vital exponent of the fractional diffusion operator.

The main findings of the present work are as follows: (i) The EEG time series at each electrode on the scalp display long memory, hence power-law scaling, as indicated by their spectral slopes in the low-frequency range. The extent of this long memory is significantly reduced from the alert state to the fatigue state. (ii) The diffusion of the EEG random field is non-Gaussian, reflecting the heterogeneity of the brain cortex. This non-Gaussianity is more severe for the alert state than the fatigue state. (iii) The system response to the initial condition, as realised by the delay parameter in the SDDE, is also stronger for the alert state than the fatigue state. The results of (ii) and (iii) in the global (whole brain) context corroborate those of (i) in the time series (individual electrode) context. The findings demonstrate the usefulness of global SDDE modelling complementing the time series approach for EEG analysis.

We will consider the cortical surface as the unit sphere 𝕊^2^, where we distribute the 32 EEG channels. The measurement of EEG at time *t* and at location **x** of an EEG channel on 𝕊^2^ is denoted by *u*(*t*, **x**). We will model the evolution of *u*(*t*, **x**), hence of the whole cortical surface, by the stochastic differential equation

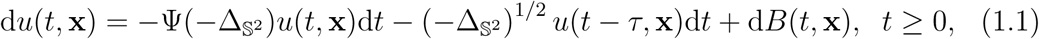

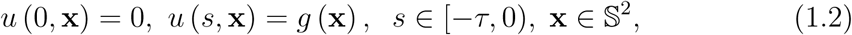

where *τ* is the delay parameter, *B*(*t*, **x**) is an *L*_2_ (𝕊^2^)-valued Brownian motion. This Brownian motion and the fractional operator

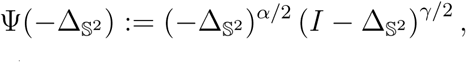

which includes 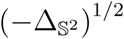 as a special case, will be defined in the next section. Briefly, 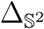 is the Laplace-Beltrami operator, which models standard diffusion on the unit sphere 𝕊^2^. The fractional operator 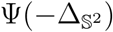 is formulated to reflect the spatial heterogeneity of the cortex and the asymptotic temporal correlation of the EEG random field. The delay parameter *τ* depicts the short-range memory and oscillation in the evolution of the system.

We will elaborate on these intrinsic features of the model in the next section. We obtain the analytical solution of the model in Section 3. This solution is presented in the form of a Karhunen-Loève expansion. This expansion allows to derive a closed-form expression for the covariance function of the random field *u*(*t*, **x**) defined by model (1.1) and (1.2). A method to estimate the key parameters of the model and the corresponding numerical schemes are then devised in Section 4. In Section 5, real EEG data on driver fatigue at all 32 channels measured on up to 50 participants will be used to estimate the fractional diffusion parameters *α, γ* and the delay parameter *τ*. Interpretation of the results is given by comparing and contrasting the alert and fatigue states of the brain. Elaboration of the above findings is also affected in this section.

## 2. Formulation of the model

In this section, we describe the components of a stochastic delay-differential equation (SDDE) to model the EEG random field on the sphere. These components include the delay response, fractional diffusion, driving noise and initial condition.

### 2.1. Random field on the unit sphere

The forward problem of EEG involves the solution of the equation **b** = *A***x**, where **b** is a vector containing information on the measured EEG field and **x** is a vector containing information on the source in the cortex generating the EEG field. *A* is an *m × n* matrix where *m* is the number of electrodes and *m* is the number of potentials to be solved on the cortical surface. Construction of the potential field from the measured EEG field is the inverse problem, which requires inverting the matrix *A*. The problem is ill-posed, and regularisation is needed. A well-known approach to obtain the inverse solution is via singular value decomposition, as discussed in [33], for example. This study examined the effects of measurement noise and the number of electrodes on the accuracy of the inverse cortical EEG solution. The authors used the spherical head model, where the cortical surface was, therefore modelled as a sphere. They found that the results obtained with a spherical head model are comparable to realistic geometry as long as the distance between the cortical surface and the scalp is similar.

In this paper, we will also assume the spherical head setting, and consider the cortical surface as the unit sphere 𝕊^2^. Each EEG channel occupies a location **x** on 𝕊^2^. The EEG measurement at a time *t* and at a location **x** is denoted *u* (*t*, **x**). As **x** varies over the entire 𝕊^2^, the field *u*(*t*, **x**) will describe the evolution of the entire cortical surface, hence representing global brain wave activity. At each time point *t, u*(*t*, **x**) is a random field on 𝕊^2^; therefore we will model its evolution by a stochastic differential equation on 𝕊^2^ as formulated in (1.1).

Let ℝ^3^ be the real 3-dimensional Euclidean space with the inner product **x** *·* **y** for **x, y** *∈*ℝ^3^ and the Euclidean norm 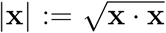. Let 𝕊^2^ := {**x** ∈ℝ^3^ : |**x**| = 1} denote the unit sphere in ℝ^3^. The sphere 𝕊^2^ forms a compact metric space with the geodesic distance dist(**x**, *y*) := arccos(**x** *y*) for **x, y** ∈𝕊^2^.

Let *L*_2_ (𝕊^2^) = *L*_2_(𝕊^2^, *µ*) be the *L*_2_-space of all real-valued square-integrable functions with respect to the Riemann surface measure *µ* on 𝕊^2^ with inner product

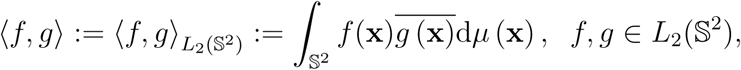

and *L*_2_-norm 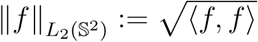.

In this paper, we use the complex-valued spherical harmonics

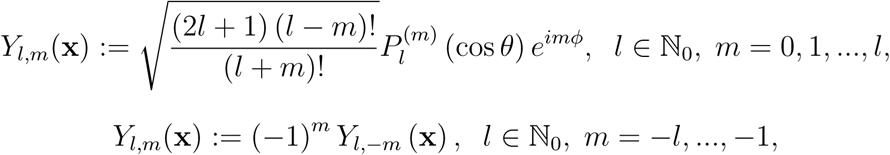

given in terms of the spherical coordinates (*θ, ϕ*) and the associated Legendre polynomial 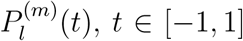, of degree *l* and order *m*. The set {*Y*_*l,m*_ : *l ∈* ℕ_0_, *m* = −*l, …, l*} is an orthonormal basis for the space *L*_2_ (𝕊^2^). For *l ≥* 0, the basis *Y*_*l,m*_ and the Legendre polynomial *P*_*l*_(**x** *·* **y**) satisfy the *addition theorem*

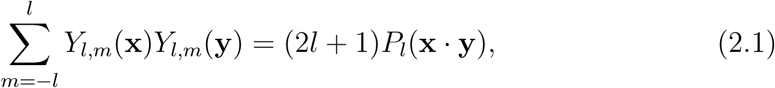

see [19, Chapter 5].

For **x** *∈𝕊*^2^, using spherical coordinates, **x** := (sin *θ* sin *ϕ*, sin *θ* cos *ϕ*, cos *θ*), *θ ∈* [0, *π*], *ϕ ∈* [0, 2*π*). The Laplace-Beltrami operator on 𝕊^2^ at **x** is given by

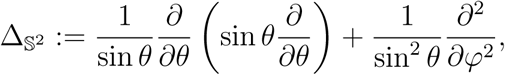

see [28, p. 38] and [13, Eq. 1.6.8]. This operator has *Y*_*l,m*_, *l ≥* 0, *m* = −*l, …, l*, as eigenfunctions with corresponding eigenvalues

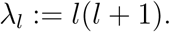

The Fourier coefficients for *f* in *L*_2_ (𝕊^2^) are given by

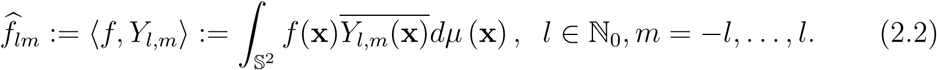

ince 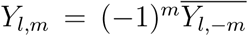 and 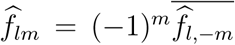 a function *f ∈ L*_2_ (𝕊^2^) has the representation

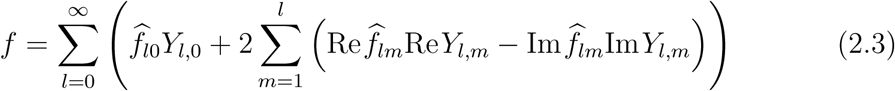

in the *L*_2_ (𝕊^2^) sense.

Given a probability space (Ω, *ℱ, P*), we denote by *L*_2_ (Ω, *P*) the *L*_2_-space on Ω with respect to the probability measure *P*, endowed with the norm 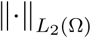. For two random variables *X, Y* on (Ω,, *ℱ, P*), we write E*X* for the expected value of *X*, Cov (*X, Y*) := E (*X* − E*X*) (*Y* − E*Y*) the covariance between *X* and *Y* and Var*X* := Cov (*X, X*) the variance of *X*.

Let *L*_2_ (Ω *×𝕊*^2^) := *L*_2_ (Ω *×𝕊*^2^, *P ⊗ µ*) be the real-valued *L*_2_-space on the product space of Ω and 𝕊^2^, where *P ⊗ µ* is the corresponding product measure. Let ℬ(𝕊^2^) denote the Borel *σ*-algebra on 𝕊^2^ and *SO*(3), the rotation group on ℝ^3^. An *ℱ⊗* ℬ (𝕊^2^)-measurable function *X* : Ω *×𝕊*^2^ *→*ℝ is called a real-valued *random field* on the sphere 𝕊^2^. We will write *X*(**x**) or *X*(*ω*) for *X*(*ω*, **x**) for simplicity if no confusion arises. We say a random field *X* is *strongly isotropic* if for any *k ∈* 𝕊, any *k* points **x**_1_, *· · ·*, **x**_*k*_ *∈ℕ*^2^ and any rotation *ρ ∈ SO*(3), the joint distributions of *X*(**x**_1_), *…, X*(**x**_*k*_) and *X*(*ρ***x**_1_), *…, X*(*ρ***x**_*k*_) coinside. We say *X* is a *Gaussian random field* on 𝕊^2^ if for each *k ∈* N and **x**_1_, *…*, **x**_*k*_ *∈ 𝕊*^2^, the random vector (*X*(**x**_1_), *…, X*(**x**_*k*_)) has a multivariate Gaussian distribution.

### 2.2. Delay response

We consider the continuum limit of a large cluster of channels on the cortical surface. A physical system with memory on the cortical surface can be represented by a Volterra-type evolution equation

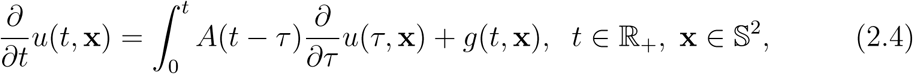

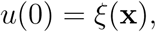

in a Banach space *E* of functions *u*(*t*, **x**) on the unit sphere 𝕊^2^. The family of linear, possibly unbounded, operators {*A*(*t*)} _*t≥*0_ can be thought of as representing the response of the system under the influence of the medium, the initial condition *ξ*(**x**) and the driving force *g* (*t*, **x**).

An approximation of Eq. (2.4) is given by

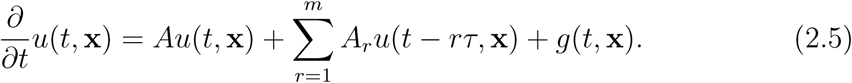

To provide a motivation for this approximation, let us consider the following situation. For *a ∈*ℝ, let *AC*_*p*_ (*a, b, E*) be the vector space of all functions *f* : [*a, b*] *→ E* which are differentiable almost everywhere on (*a, b*) with derivative in *L*_*p*_ (*a, b*; *E*), and such that

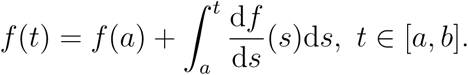

The functional

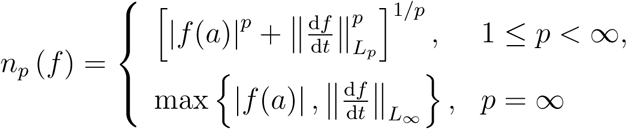

defined in [15] is a norm on *AC*_*p*_ (*a, b, E*), which is then a Banach space isometrically isomorphic to *E × L*_*p*_(*a, b*; *E*).

For *f ∈ AC*_*p*_(*a, b, E*), assuming *A*(*t*) is of scalar type on each interval [*t*−*τ, t*], [*t*− 2*τ, t* − *τ*], *…*,

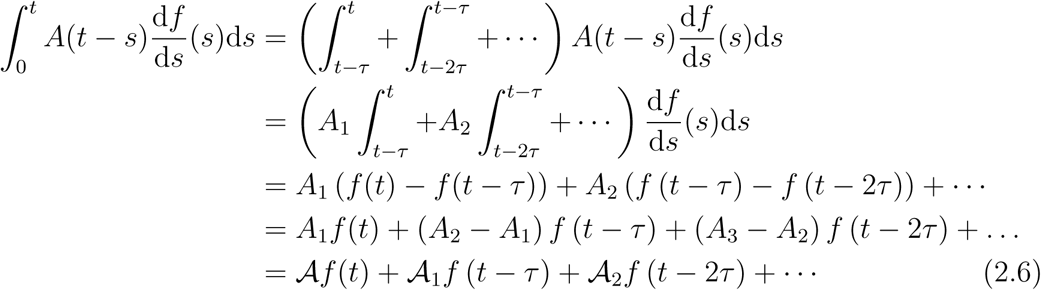

Note that an operator *A*(*t*) is of scalar type if *A* (*t*) = *a*(*t*)*A*, where *A* is a closed linear, densely defined operator in *E*, and 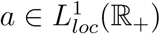, a scalar kernel. In (2.6), we assume *A*(*t*) is of scalar type with *a* (*t*) = 1 on each interval [*t* −*τ, t*], [*t* −2*τ, t* −*τ*], *…*. Truncation of the expansion (2.6) at *mτ* then yields the approximation (2.5).

### 2.3. Fractional diffusion operator

In this subsection, we introduce the fractional diffusion operator. Let *α >* 0, *γ ≥*0, and

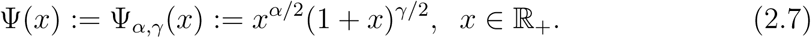

The fractional diffusion operator 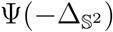 used in the model (1.1) is defined as

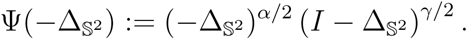

Using (2.7), the operator 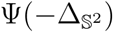 has eigenvalues

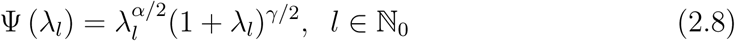

(see [14, p. 119-120]), and

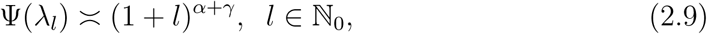

where *a*_*l*_ *≍b*_*l*_ means *cb*_*l*_ *≤ a*_*l*_ *≤ c′ b*_*l*_ for some positive constants *c* and *c*^*′*^.

The operator 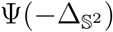 on 𝕊^2^ is the counterpart of the fractional diffusion operator in R^*n*^. We recall that the operator 𝒜:= − (−Δ)^*α/*2^(*I* −Δ)^*γ/*2^, which is the inverse of the composition of the Riesz potential (−Δ)^−*α/*2^, *α ∈* (0, 2], defined by the kernel

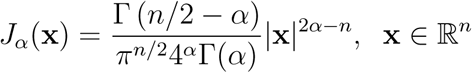

and the Bessel potential (*I* − Δ)^−*γ/*2^, *γ ≥* 0, defined by the kernel

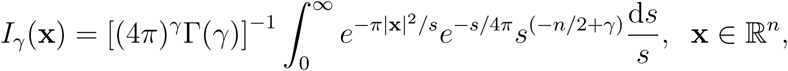

(see [34]) is the infinitesimal generator of a strongly continuous bounded holomorphic semigroup of angle *π/*2 on *L*_*p*_(ℝ^*n*^) for *α >* 0, *α* + *γ ≥* 0 and any *p ≥* 1, as shown in [5]. This semigroup defines the Riesz-Bessel distribution if and only if *α ∈* (0, 2], *α* + *γ ∈* [0, 2]. Let *X* (*t*) denote the process, named the Riesz-Bessel motion (see [3, 4]), defined by this Riesz-Bessel distribution. When *γ* = 0, the fractional Laplacian − (−Δ)^*α/*2^, *α ∈* (0, 2], generates the Lévy *α*-stable distribution. As *t →∞, t*^−1*/α*^*X* (*t*) converges in distribution to a symmetric *α*-stable random variable, while as *t →*0, assuming *α* + *γ >* 0, *t*^−1*/*(*α*+*γ*)^*X* (*t*) converges in distribution to a symmetric (*α* + *γ*) −stable random variable [5]. Consequently, the exponent *α* controls the tail behaviour of the distribution of the Riesz-Bessel motion and indicates how often large jumps occur, while the exponent *γ*, through the value of the sum *α* + *γ*, controls the small-scale structure and describes the multifractal behaviour of the Riesz-Bessel motion. These results are used to interpret the meaning of the component 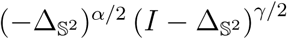 when we fit the EEG data for the alert and fatigue states in Section 5.

#### 2.3.1 Fractional diffusion in a heterogeneous medium

Let *µ*_*M*_ be a finite Borel measure with compact support *M ⊂*ℝ^*n*^. We say that the measure *µ*_*M*_ is a *d*(*·*)-measure if, for every **x** *∈ M*, it satisfies

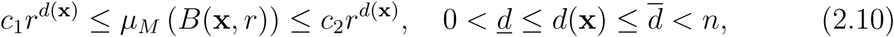

for *r ∈*(0, *r*_0_) for some fixed *r*_0_, where *c*_1_, *c*_2_ are positive constants, and *B*(**x**, *r*) denotes the closed ball with center **x** and radius *r*. The exponent *d*(**x**) is the local dimension of *µ*_*M*_. If *d*(**x**) = *d*, (*µ* − a.e.), *d* is called the fractal dimension of *µ*_*M*_. [32] showed that the transition probability densities of a class of processes can be constructed from the fundamental solution of the equation

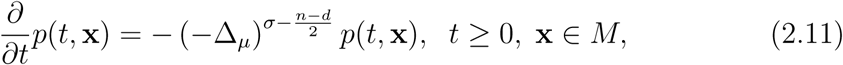

where *M* is a compact set in ℝ^*n*^ whose Borel measure *µ* has fractal dimension *d*, and (−Δ_*µ*_)^*s*^ is the negative Laplacian on *M*. In (2.11), *σ* is the regularity order of a Markov diffusion. The exponent 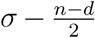 shows that this regularity order is reduced by the amount 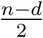, which is the fractal effect due to the fractal domain *M* in ℝ^*n*^. This effect is built into the exponents of the diffusion operator in our model (1.1).

#### 2.3.2 Asymtotic temporal correlation of fractional diffusion

To understand the temporal correlation of the fractional diffusion of the EEG field *u*(*t*, **x**), let us look at a simpler version of model (1.1) without the delay term, written in the form of stochastic partial differential equation

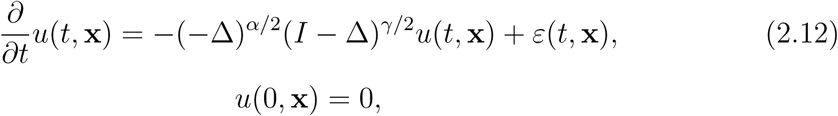

where **x** now varies in the two-dimensional planar cortical sheet ℝ^2^, and *ε*(*t*, **x**) is Gaussian space-time white noise (defined as a random Schwartz distribution). In this situation of an unbounded spatial domain, Fourier transform techniques can be applied, and the spectral density of the solution of (2.12) can be derived when the solution is stationary. This derivation clarifies the meaning of the parameters *α* and *γ*. In fact, for *t ∈*ℝ_+_, we denote by 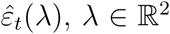, the complex-valued generalised random function defined by the following weak-sense identity in *L*^2^(ℝ_+_ *×*ℝ^2^) :

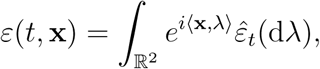

with 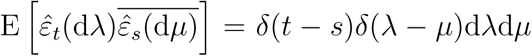 for all *λ, µ ∈*ℝ^2^ and *t, s ∈*ℝ_+_. As established in [2],

##### Proposition 2.1.

*A real-valued zero-mean solution, in the mean square sense, of* (2.12) *defined on* ℝ_+_ *×* ℝ^2^, *under zero initial condition and assuming α* + *γ >* 2, *is given by*

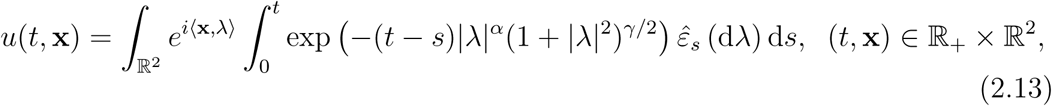

*where the integrals are interpreted in the mean-square sense. In addition, if α <* 2, *the process is asymptotically stationary with its asymptotic temporal covariance function given by*

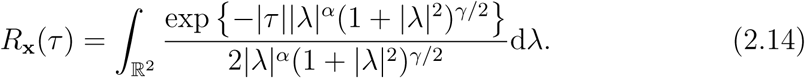

Changing (2.14) to polar coordinates yields

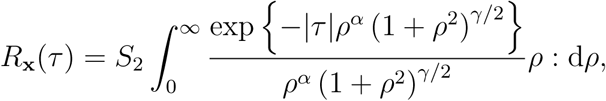

where *S*_2_ is a constant resulting from the change to polar coordinates. Making the change of variable *u* = |*τ*|*ρ*^*α*^, we have

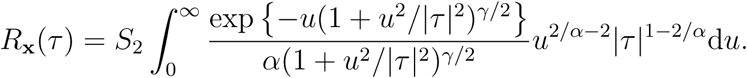

From the dominated covergence theorem,

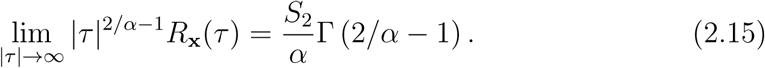

Thus, we have *asymptotic temporal correlation*, which is governed by the exponent *α*. It is seen from (2.14) that the covariance function has a slower decay than an exponential function and, if *α >* 1, the temporal process will exhibit long-range dependence. This finding is significant in the sense that we observe temporal long-range dependence, with 1 *< α <* 2, even in the case of the (infinite-dimensional) Ornstein-Uhlenbeck process (2.12). This result is a useful tool in our investigation of existence of long-range dependence, hence power-law scaling, of global brain activity.

### 2.4. Driving noise

A real-valued Brownian motion *β*(*t*), *t ≥* 0, with variance *σ*^2^ at *t* = 1 is a centered Gaussian process on ℝ_+_ which satisfies

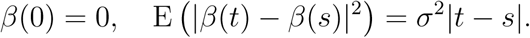

The variance of *β*(*t*) is then E (|*β*(*t*)|^2^) = *σ*^2^*t, t >* 0.

#### Definition 2.2.

*Let β*^(1)^(*t*) *and β*^(2)^(*t*) *be independent real-valued Brownian motions with variance* 1 *(at t* = 1*). A complex-valued Brownian motion B*(*t*), *t ≥* 0, *with variance σ*^2^ *is defined as*

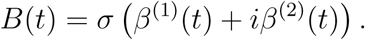

The noise in Eq. (1.1) is modelled by an *L*_2_(𝕊^2^)-valued Brownian motion *B*(*t*) defined as follows.

#### Definition 2.3.

*Let b*_*l*_ *>* 0, *l ∈*ℕ_0_ *satisfy* 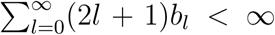. *Let B*_*lm*_(*t*), *t* 0, *l ≥* ℕ_0_, *m* = −*l, …, l be a sequence of independent complex-valued Brownian motions on* ℝ_+_ *with variance b*_*l*_ *at t* = 1 *and* Im*B*_*l*0_(*t*) = 0 *for l ∈* ℕ_0_, *t ≥* 0. *For t ≥* 0, *the L*_2_(𝕊^2^)*-valued Brownian motion is defined by the following expansion (in the L*_2_ (Ω *×𝕊*^2^) *sense) in spherical harmonics with Brownian motions B*_*l,m*_(*t*) *as coefficients:*

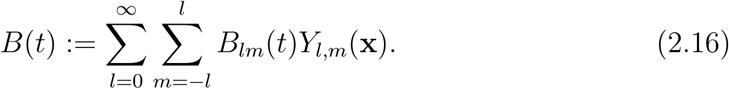

We also call *B*(*t*) in (2.16) a Brownian motion on 𝕊^2^, written *B*(*t*, **x**), **x** *∈𝕊*^2^. The random field *B*(*t*) in (2.16) is well-defined since for *t ≥* 0, by Parseval’s identity,

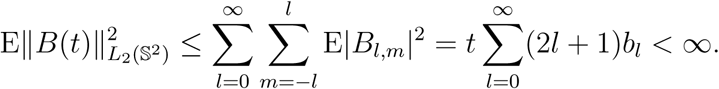

In this paper, we let *B*(*t*) be real-valued. For *l ∈* ℕ_0_, let

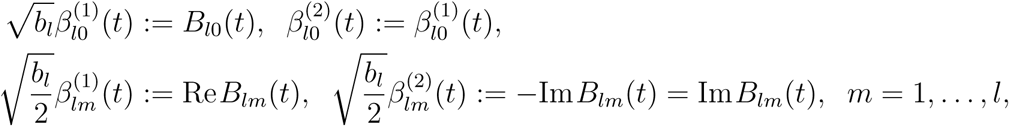

in law. Then, 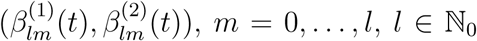, is a sequence of independent real-valued Brownian motions with variance 1 (at *t* = 1). In view of (2.3), we can write *B*(*t*) for *t ≥* 0, in the *L*_2_(Ω *×𝕊*^2^) sense, as

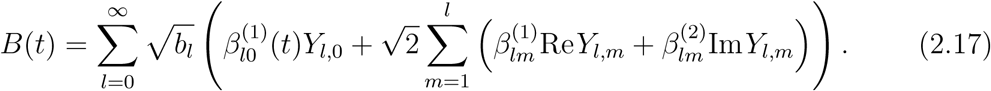

For a bounded measurable function *g* on ℝ_+_ (which is deterministic), the stochastic integral 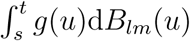 can be defined as a Riemann-Stieltjes integral. The *L* (𝕊^2^)-valued stochastic integral 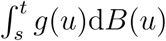, for *t > s ≥* 0, can then be defined as an expansion in spherical harmonics with coefficients 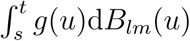 as

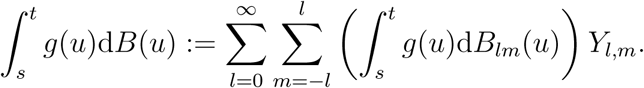

### 2.5. Initial conditions

Eq. (1.1) is solved under the initial conditions *u*(0, **x**) = 0, *u*(*s*, **x**) = *g*(**x**), *s ∈* [−*τ*, 0), **x** *∈𝕊*^2^, where we assume *g*(**x**) to be a strongly isotropic Gaussian random field on 𝕊^2^. It has the expansion

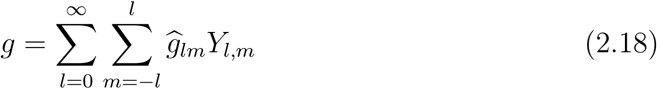

in the *L*_2_(Ω *×𝕊*^2^) sense, where the Fourier coefficients 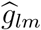 are independent and normally distributed. We assume 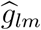 has mean 0 and variance *c*_*l*_.

## 3. Analytical solution

In this section, we show the analytic solution of the proposed SDDE by Karhunen-Loève expansion and prove its convergence. The solution relies on the theory of fundamental solution of delay-differential equations in Banach spaces outlined in the Appendix. We then provide an explicit formula for the covariance function of the solution.

### 3.1. Karhunen-Loève representation

Model (1.1) is a special case of (A.1) with 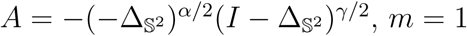 and 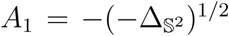. As noted in Subsection 5.3.1, the operator 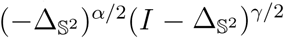 has eigenvalues 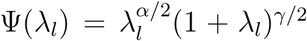, with *λ*_*l*_ = *l*(*l* + 1), *l ∈* ℕ_0_, and eigenfunctions {*Y*_*l,m*_ : *l ∈* ℕ_0_, *m* = −*l, …, l*}. Thus,

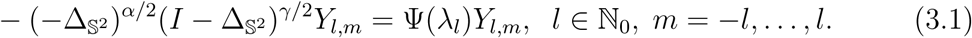

Therefore, the semigroup generated by 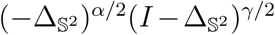 has the representation

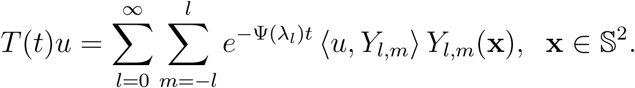

It is seen that 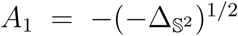 satisfies condition **H**_3_; thus, using (A.6), the fundamental solution *G*(*t*) of (1.1) is given by, for *t ∈* [(*k* − 1)*τ, kτ*] and 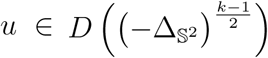,

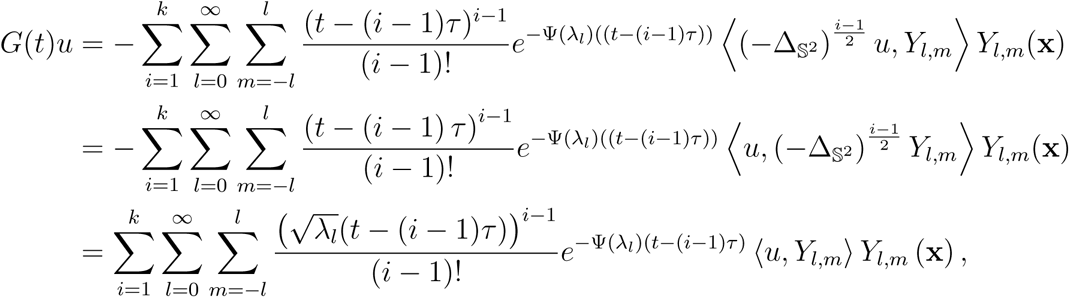

where the last equality is due to that 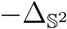 is essentially self-adjoint [24, p. 299], and (3.1) is used for *γ* = 0 and *α* = *i* − 1. We then obtain

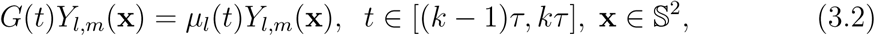

with

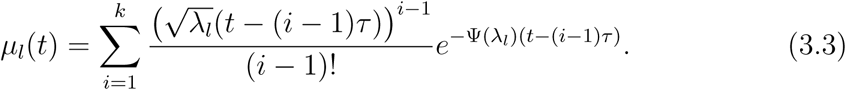

#### 3.1.1. Periodic motion generated by the delay operator

Now let us look at the effect of the delay in the SDE. Denote by *𝒞* = *C* ([−*τ*, 0]; *E*) the Banch space of continuous maps *ψ* : [−*τ*, 0] *→ E* with the sup norm. We consider the governing component of Eq. (1.1) in the phase space *𝒞*:

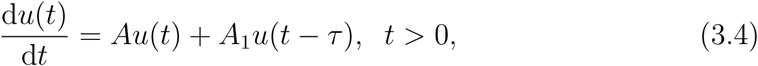

where the operators *A, A*_1_ are as defined above. Let **C** =*C* ([−*τ*, 0]; ℝ). For each *l ∈* ℕ_0_, we define the maps *A*_*l,m*_ : **C** *→*ℝ by

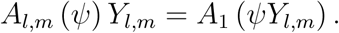

We introduce the subspaces

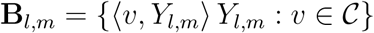

of *𝒞* which satisfy *A*_1_**B**_*l,m*_ *⊂* span{*Y*_*l,m*_}. Then, on **B**_*l,m*_, the linear equation (3.4) is equivalent to the functional delay differential equation on ℝ :

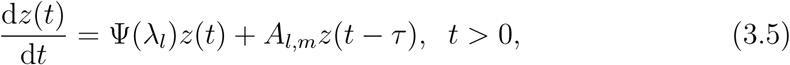

(see [17, Eq. 1.6_k_]) with characteristic equations

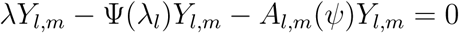

(see [17, Eq. 1.3_k_]), which reduce to

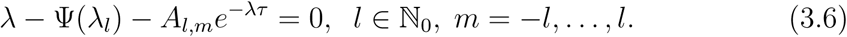

For each *l*, we re-write Eq. (3.6) as

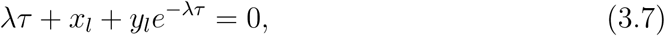

where *x*_*l*_ = −Ψ(*λ*_*l*_)*τ, y*_*l*_ = −*A*_*l,m*_*τ*. We look for solutions to Eq. (3.7) of the form *λ* = *iω*. Eq. (3.7) then becomes

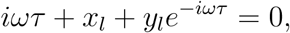

which yields, for the real and imaginary parts,

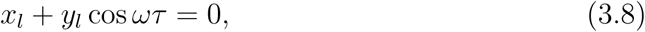

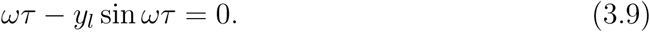

From Eq. (3.9),

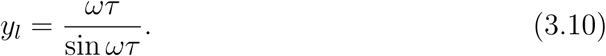

Inserting this into (3.8) yields

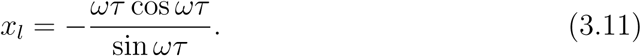

The parametric equations (3.10) and (3.11) describe a curve of *y*_*l*_ vs *x*_*l*_ in terms of *ωτ*. For a range of *ωτ*, such as 0.5*π < ωτ <* 0.9*π*, the curve is almost linear, which is known as the Hopf curve. We can re-write Eq. (3.5) in terms of *x*_*l*_ and *y*_*l*_:

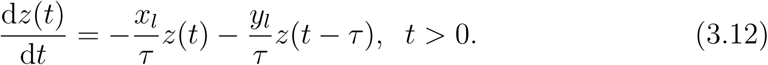

An equation of the form (3.12) was used to study periodic breathing in [18]. In fact, Eq. 2 in [18] is of the form (3.12), and their characteristic equation is of the form (3.6). By numerical simulation, it was found that, for the values of (*x, y*) below the Hopf curve, the equilibrium solution *z*(*t*) = 0 is stable, while above the curve it is unstable and there are periodic oscillations. That is, periodic motion occurs for (*x, y*) above the Hopf curve as demonstrated in Fig. 3 in [18].

**Figure 1:**
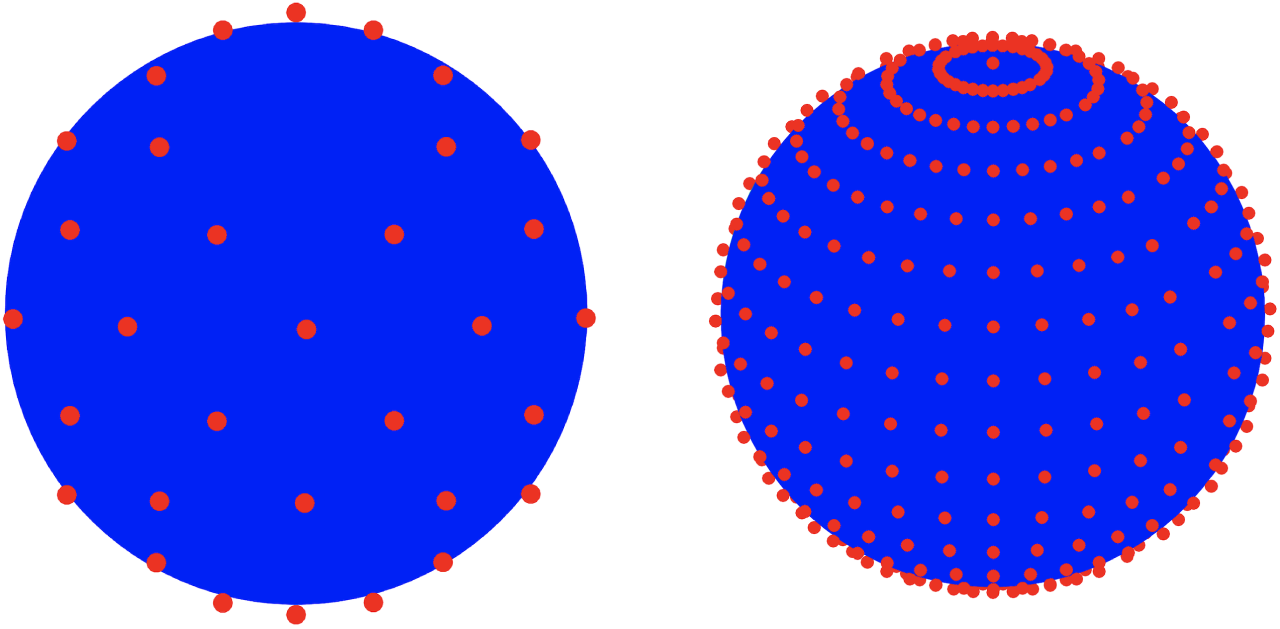
Left: Position of 32 channels for EEG signals. Right: 512 nodes of Gauss-Legendre product rule on the sphere

**Figure 2:**
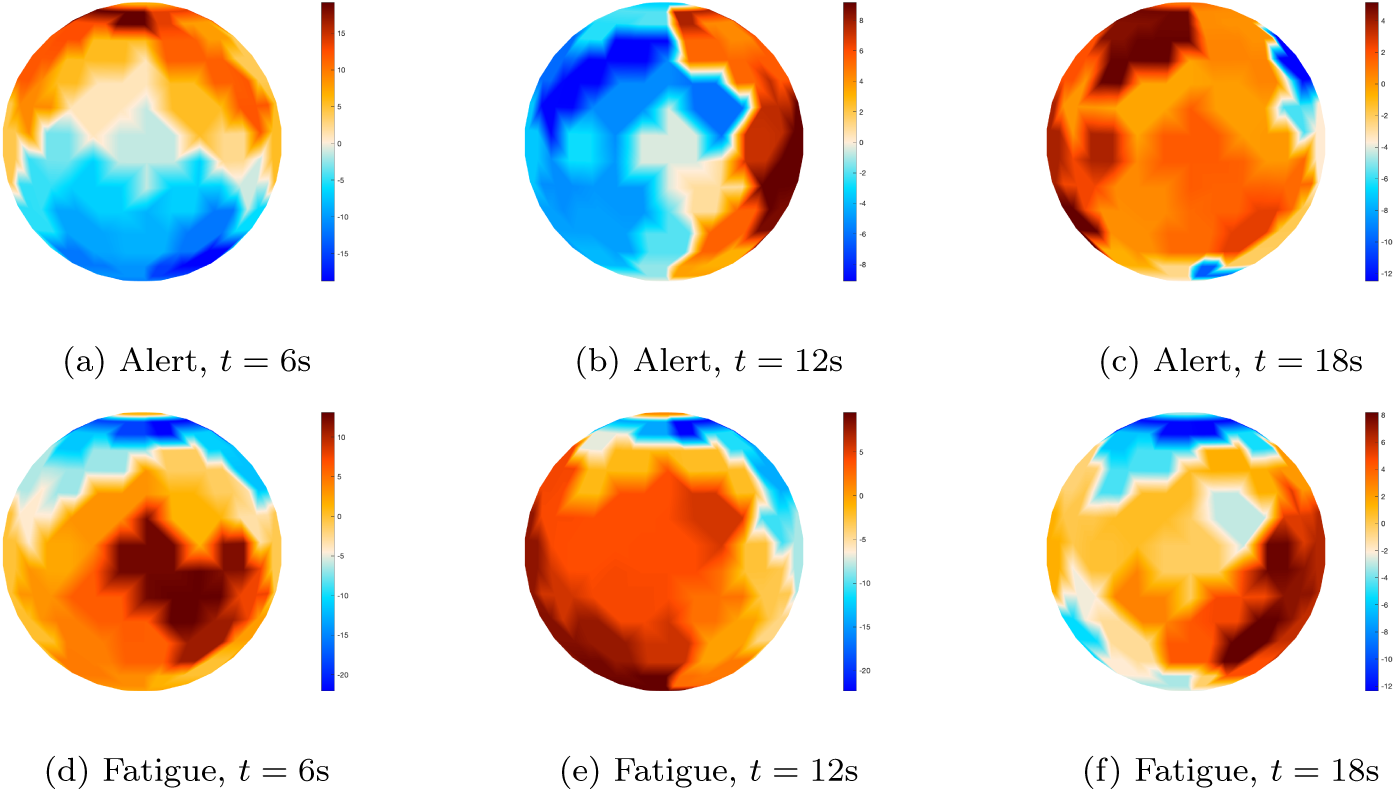
Two states of brain wave activity at *t* = 6, 12, 18 seconds

**Figure 3:**
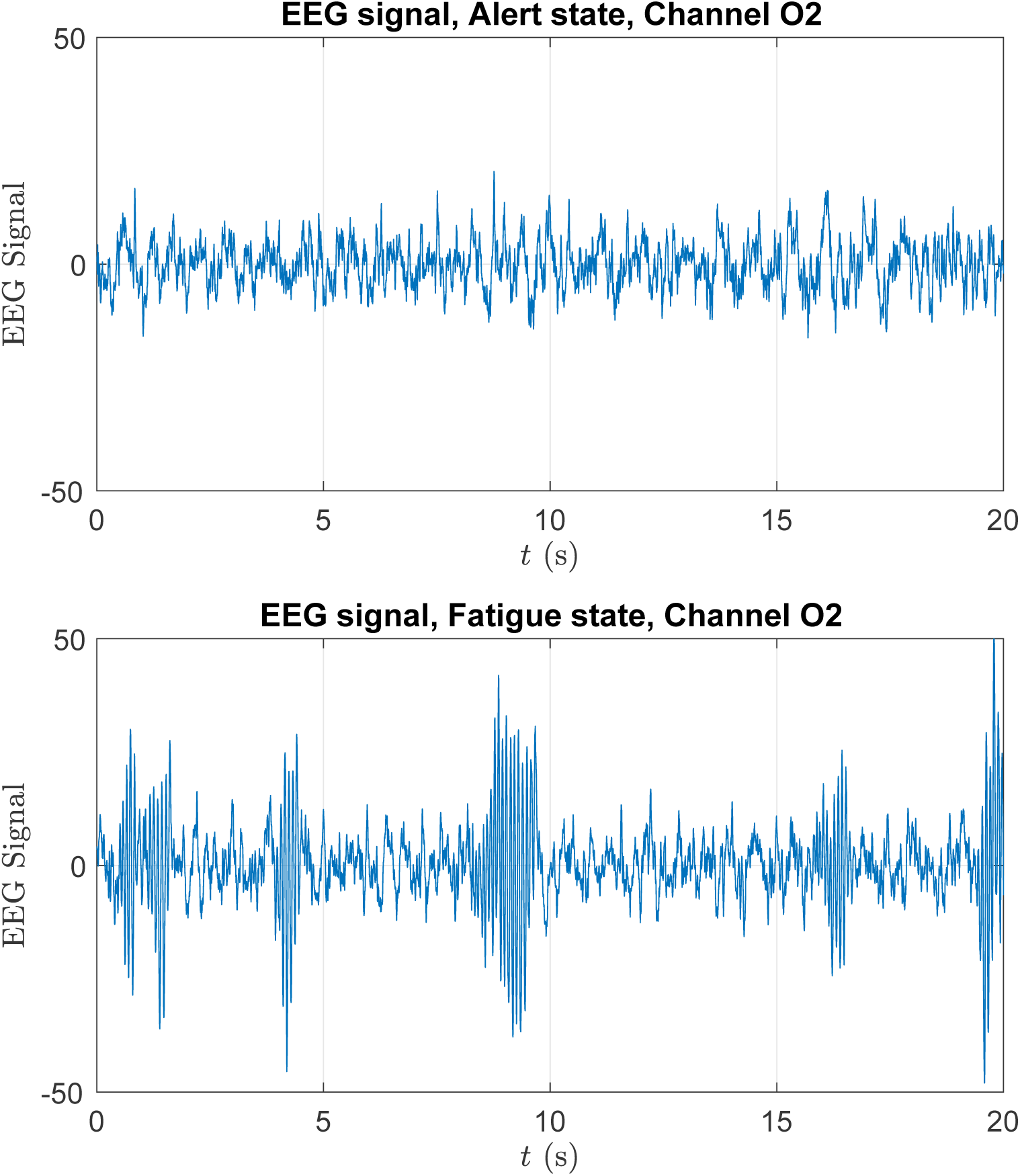
EEG time series at channel O2

Going back to model (1.1) and (1.2), continuing from (3.3) and using (A.7), its mild solution is then given by

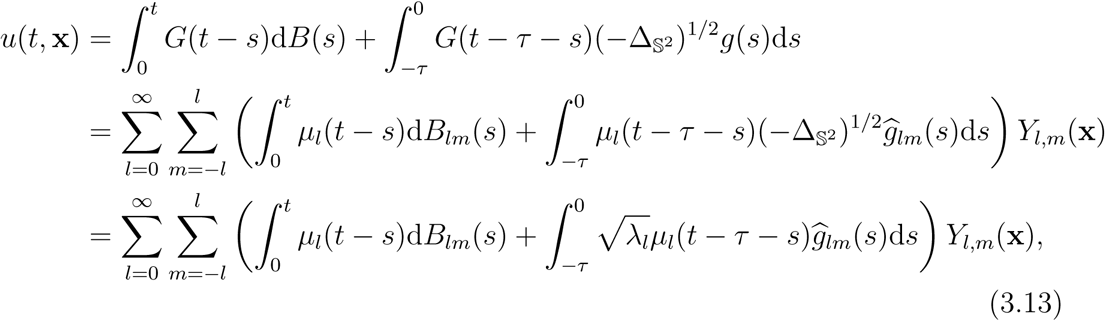

where the second equality uses 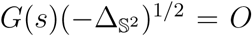 if *s <* 0, and the last equality uses (2.16), (2.18), (3.2) and the commutativity of *G*(*t*) and 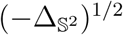, and (3.1) is used when *γ* = 0, *α* = 1. Noting the condition *u*(*s*, **x**) = *g*(**x**), *s ∈* [−*τ*, 0), **x** *∈𝕊*^2^ in (1.2), the Karhunen-Loève representation of the mild solution of model (1.1) and (1.2) is then

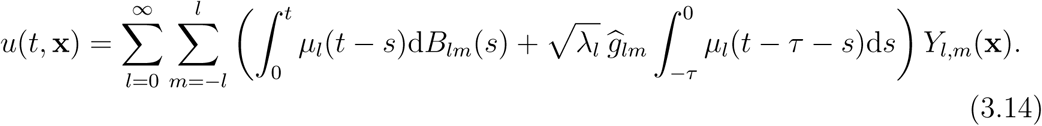

##### Theorem 3.1.

*The expansion of u*(*t*, **x**) *in* (3.14) *converges in L*_2_ (Ω *×𝕊*^2^). *Proof*. By Parseval’s identity, the squared *L*_2_-norm of *u*(*t*) at *t* is

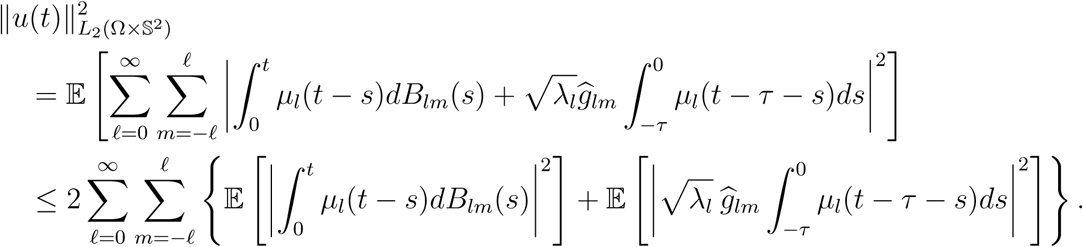

In view of Itô’s isometry theorem [31, Lemma 3.1.5],

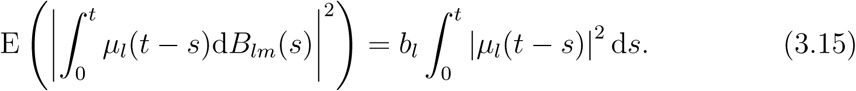

Then

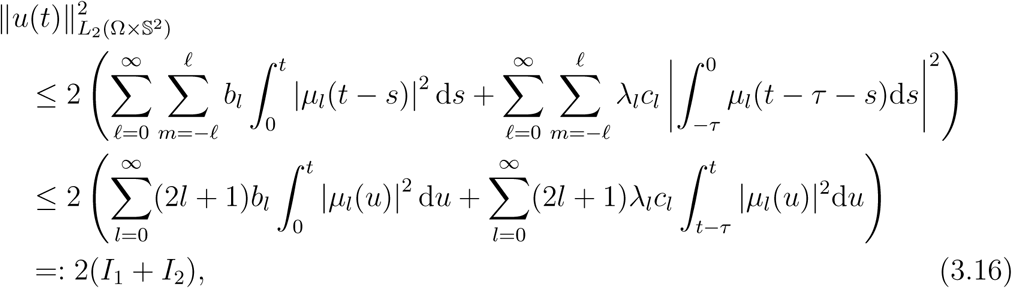

where *c*_*l*_ is the variance of 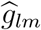. For the first term of (3.16), by (3.3) and Jensen’s inequality,

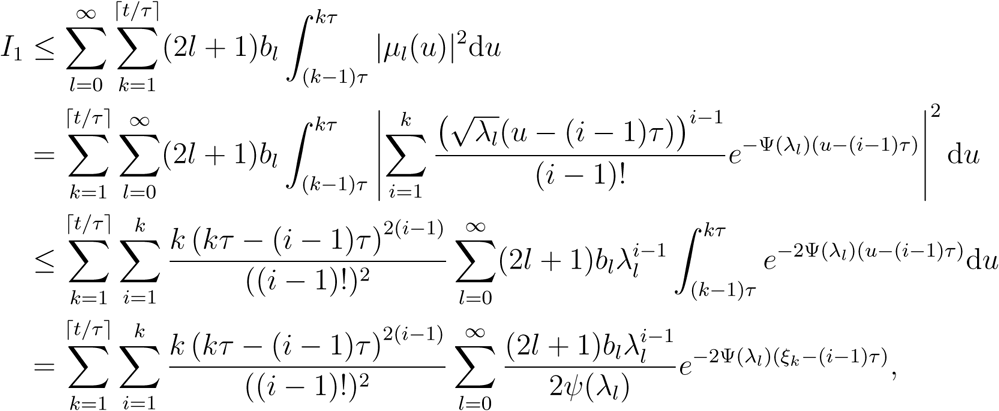

where the last line holds for some *ξ*_*k*_ in ((*k*− 1)*τ, kτ*) by the mean value theorem. As *ξ_k_ >* (*k* − 1)*τ ≥* (*i* − 1)*τ*, the series for 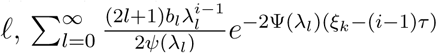 converges for *i* = 1, *…, k* and *k* = 1, *…, ⌈t/τ⌉*. Thus, *I*_1_ *< ∞*.

In a similar way, for some 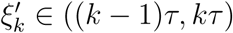,

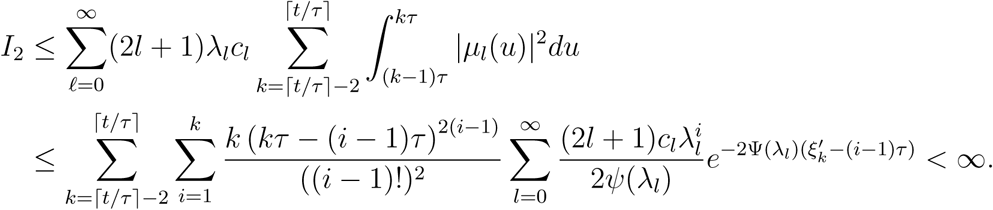

This together with *I*_1_ *< ∞* and (3.16) shows 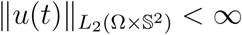.

□

### 3.2. Covariance function

As given by (3.13), the solution *u*(*t*, **x**) has mean 0 by assumption. At a fixed time *t*, its covariance function is then

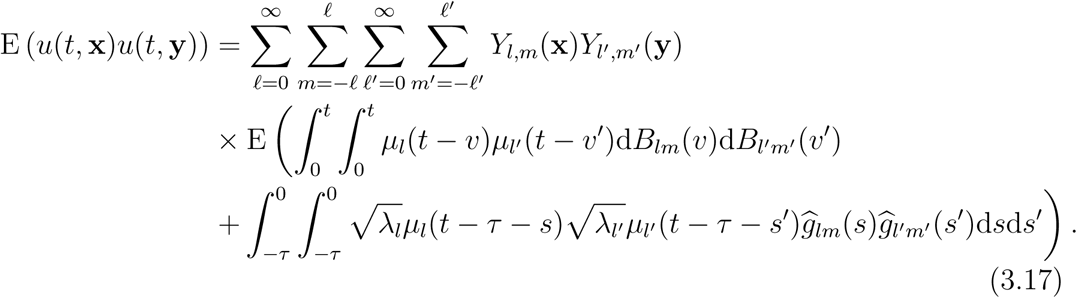

By the independence of *B*(*t*) and *g*(*t*), and the independence of the coefficients at different indices (*l, m*), the covariance function (3.17) becomes

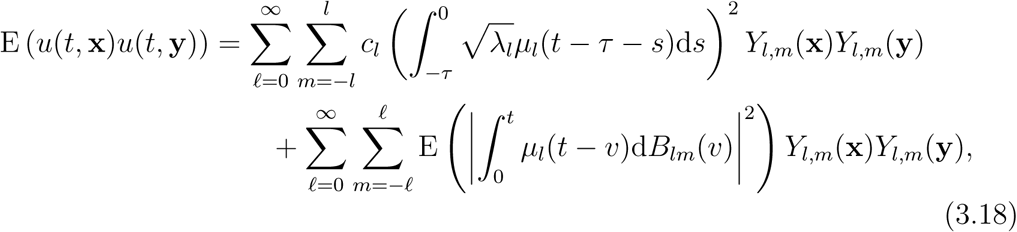

where we used (2.18). Thus, using (3.15) and the addition theorem (2.1) again, Eq. (3.18) becomes

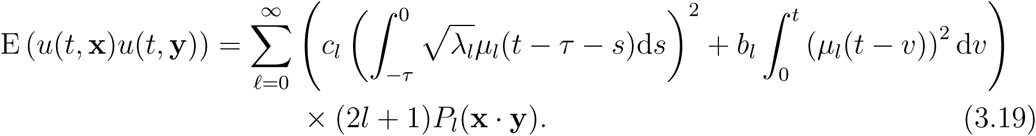

Using P_l_ (x _· x) = P_l_(1) = 1, we then obtain the variance of u(t, x) as

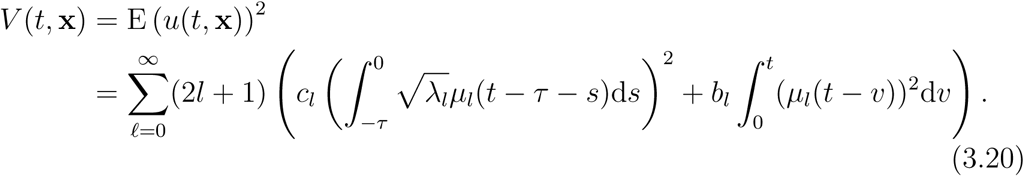

## 4. Methods for parameter estimation

In this section, we provide some methods for numerical estimation of the fractional exponents *α* and *γ*, the delay parameter *τ*, and the exponent *θ* in the variances of the initial condition of the SDDE (1.1) and (1.2).

### 4.1. Estimation of the fractional diffusion

To estimate the exponents *α* and *γ*, we let

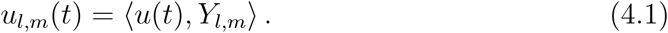

Then, using (3.14),

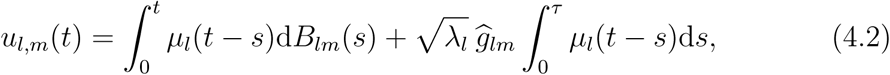

where we have used the change of variable *τ* + *s → s* in the second integral. We noted in Subsection 2.1 that, for a bounded measurable function *f* on R_+_ (which is deterministic), the stochastic integral 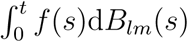 can be defined as a Riemann-Stieltjes integral.

We now consider the representation

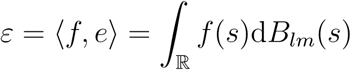

for 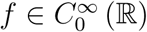, the space of infinitely differentiable functions with compact support in ℝ. The function ⟨*f, e⟩* is linear and continuous with respect to the *L*_2_-norm over 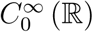. We may treat *ε* as a random Schwartz distribution, and identify *ε*(*t*) with the derivative 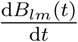. Then, Eq. (4.2) can be written formally as

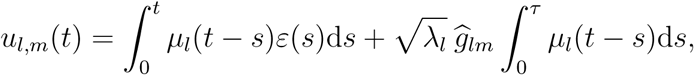

from which and (3.3) we obtain

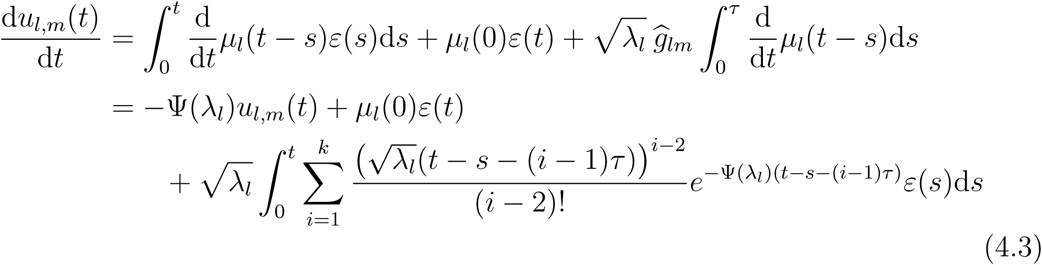

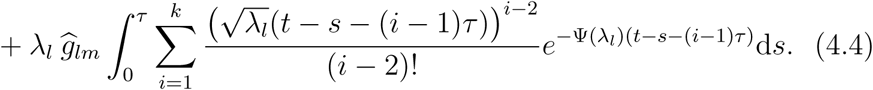

For this initial estimation of *α, γ*, we will compute the values of *u*_*l,m*_(*t*) from (4.1) for large *l* so that

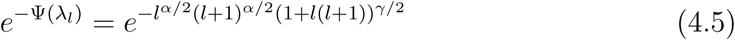

is small. In this setting, the integrals in (4.3) and (4.4) are approximately zero for *l* sufficiently large. Then, *u*_*l,m*_(*t*) satisfies the equation

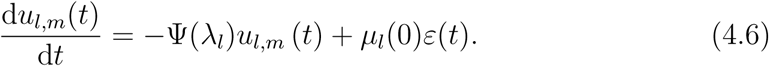

For each *l, m*, Eq. (4.6) is an Ornstein-Uhlenbeck equation for *u*_*l,m*_(*t*) driven by white noise *ε*(*t*). By [22], for each *m* = −*l, …, l*,

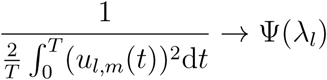

as *T → ∞*. Then, using *n* samples *u*_*l,m*_(*t*_1_), *…, u*_*l,m*_(*t*_*n*_), *t*_*j*_ = *jT/n*,

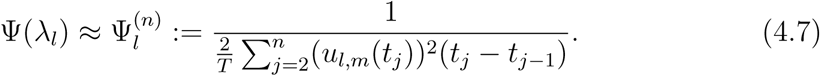

By (4.5) and (4.7), we propose to estimate the parameters *α* and *γ* by solving the following nonlinear least squares problem: for *L*_2_ *≥L*_1_ *≥*1 and *L*_1_, *L*_2_ sufficiently large,

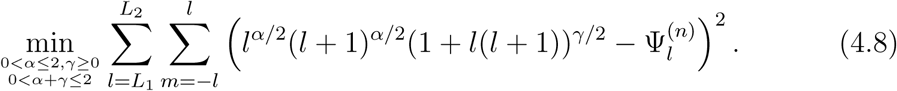

When *L*_1_ is sufficiently large, Eq. (4.8) is approximately by

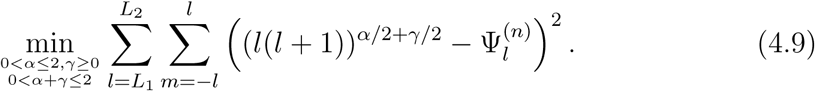

In practice, we first estimate *α* + *γ* for *L*_1_ sufficiently large by (4.9), then estimate *α* using a smaller *L*_1_ and setting *γ* = 0 in (4.8).

### 4.2. Estimation of the delay parameter and initial condition

In this subsection, we estimate the parameters for the delay operator and the initial condition. This estimation is proceeded under the assumption that the delay is the response of the system due to the initial condition *u* (*s*, **x**) = *g* (**x**) for *s ∈* [−*τ*, 0) and **x** *∈𝕊*^2^. We assume the variances of 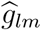 take the form

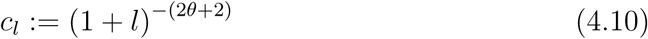

for *θ >* 1. Then, the estimation involves the parameters *τ* and *θ*.

Going back to the form (4.2) and using (3.20) we find

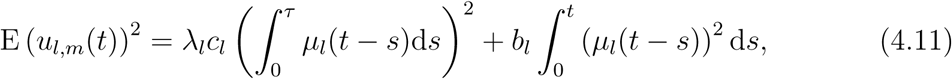

where the second term on the right-hand side vanishes as *t →* 0. We recall that *µ*_*l*_(*t*) of (3.3) is defined for *t ∈* [(*k* − 1) *τ, kτ*]. Thus, for 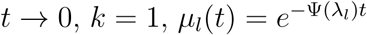 and

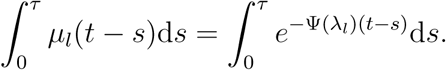

Since 0 *< t* − *s < t*, we get *t* − *s →* 0 as *t →* 0. Hence,

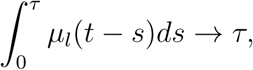

and by (4.11),

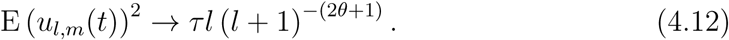

We compute E (*u*_*l,m*_(*t*))^2^ by discretising the integral with *n* sample times as

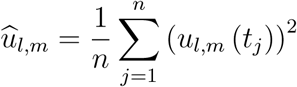

over a partition 0 = *t*_1_ *< t*_2_ *< … < t*_*n*_ = *t*_0_ of the interval [0, *t*_0_], where *t*_0_ is a small positive number. Then, the formula (4.12) suggests to estimate the parameters *τ* and *θ* by solving the nonlinear least-squares problem: for a positive integer *L*,

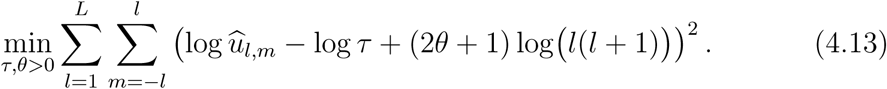

## 5. Parameter estimation on real EEG data

### 5.1. EEG dataset

We view the brain as part of the sphere 𝕊^2^ *⊂*ℝ^3^. The left panel of Fig. 1 shows the location of the 32 EEG channels. At each channel, the EEG brain wave was measured for 20 seconds composing of 5120 time points. Fig. 2 displays the EEG patterns on 𝕊^2^ at *t* = 6, 12, 18 seconds. Figures 2a,b,c show the EEG for the alert state, while Figures 2d,e,f for the fatigue state. They illustrate two apparently different patterns of the random field of brain wave activity on 𝕊^2^.

### 5.2. Fourier coefficients

We compute the Fourier coefficients *u*_*l,m*_ by the fast spherical harmonic transform [26, 25] using EEG measurements. To increase the accuracy of evaluation, we extrapolate the measured EEG data at 32 channels on 𝕊^2^ to 512 nodes of the Gauss-Legendre product rule, as shown in the right panel of Fig. 1. The Gauss-Legendre tensor product rule is a (polynomial-exact but not equal-area) quadrature rule with positive weights on 𝕊^2^ [21, 35]. The tensor product of the Gauss-Legendre zeros on [−1, 1] determines the latitude and equally-distributed nodes on the circle at a latitude. The Gauss-Legendre rule with *N* nodes is exact for polynomials of degree *n*:

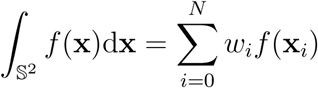

for all spherical polynomials *f* of degree up to *n*, and the number of points *N* = *n ×* (⌈(*n* − 1)*/*2⌉ + 1). The function value at an extrapolation point is set equal to the value at the closest channel.

### 5.3. Parameter estimation

In this section, we estimate the parameters of the SDDE using real EEG data. The parameters include *β, α, α* + *γ, τ* and *θ*.

#### 5.3.1. Power-law scaling of EEG time series

For a discussion on power-law scaling in EEG time series, let us recall a few facts on fractional Gaussian noise. The generalised derivative (in the sense of Schwartz distributions) of fractional Brownian motion *B*_*H*_, 0 *< H <* 1, is called fractional Gaussian noise. For *H ∈* [1*/*2, 1), this noise process is commonly known as 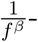-noise, with 0 *≤ β* = 2*H* − 1 *<* 1, where *f* stands for frequency. To avoid confusion, we will write 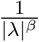 when we refer to the spectral density of 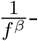 -noise. In the range 0 *< β <* 1, 1*/f* ^*β*^-noise is stationary, strongly dependent and interpolates between white noise (1*/f* ^0^-noise, which has a constant spectral density) and pink noise (1*/f* ^1^-noise, which has a spectral density proportional to the reciprocal of the frequency). For 1 *< β* = 2*H* + 1 *<* 3, 1*/f* ^*β*^-noise is non-stationary and possesses short-range dependence for 1 *< β <* 2 (0 *< H <* 1*/*2) and long-range dependence for 2 *< β <* 3 (1*/*2 *< H <* 1). The change from stationary strongly dependent 1*/f* ^*β*^-noise, 0 *< β <* 1, to nonstationary 1*/f* ^*β*^-noise, 1 *< β <* 3, can be considered as a change of states of the system. Pink noise (1*/f* ^1^-noise) represents the changing point between these two states.

As an illustration, we show the EEG time series at channels O2 and P4 in the alert and fatigue states. There is clear intermittency in the O2 time series in the fatigue state. This is depicted by a singularity at a frequency in the range 100 *< ω <* 300 in its periodogram. The appearance of this frequency for intermittency is likely due to the closed eye tendency when a driver is tired. Similar strong intermittency is found in the P4 time series in the fatigue state.

The spectral slopes of the O2 and P4 time series at low frequencies (*ω ≤* 100) are *β* = 1.33 and 1.23 respectively for the alert state, and *β* = 1.37 and 1.28 respectively for the fatigue state. These slopes are obtained via the regression of log *f* (*λ*) against log |*λ*| based on the 1*/f* ^*β*^-noise model log *f* (*λ*) ∼−*β* log |*λ*| as *λ→* 0. The estimated slopes indicate that the alert-state time series are nonstationary and possess short-range dependence with *H* = 0.165 and 0.115 respectively using the formula 2*H* + 1 = *β*. The Hurst indices are *H* = 0.185 and 0.14 for O2 and P4 respectively in the fatigue state, indicating that there is no significant change (in the low-frequency behaviour of these time series) from the alert state to the fatigue state.

#### 5.3.2. Global power-law scaling

The covariance function of (2.15) is of the form

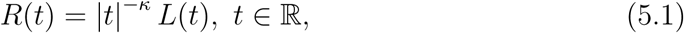

with 0 *< κ <* 1 for 1 *< α <* 2, and *L* (*x*) being a function slowly varying at infinity. A Tauberian theorem [27, p. 66] implies that the spectral density *f* (*λ*) corresponding to *R*(*t*) behaves as

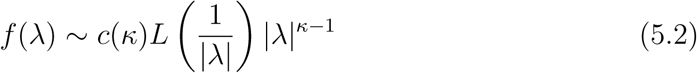

as |*λ*| *→* 0, where the Tauberian constant 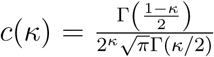. Thus, at a fixed location, the asymptotic solution of (2.12) behaves as 1*/f* ^*β*^-noise, with *β* = 2 (1 −1*/α*). Consequently, the parameter *α* of the fractional diffusion operator 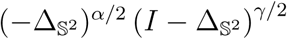 of (1.1) quantifies its 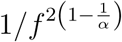-noise behaviour at low frequencies. Under the assumptions of this global model, there may not be any formula connecting this low-frequency behaviour via *β* = 2 (1 −1*/α*) with the bahaviour via *β* = 2*H* + 1 of the EEG time series investigated above because the *α*-component in the operator 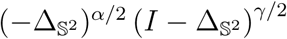 picks up only a part of the memory in the EEG global field. The remaining part is embedded in the delay operator 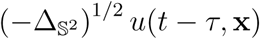. However, as described, this 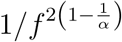-noise behaviour can be used as an indicator to distinguish between the alert and fatigue states. This interpretation is included in the next subsection.

#### 5.3.3. Behaviour of global EEG

We use (4.7) and (4.13) to estimate the parameters *α, γ* and the delay parameter *τ* as well as the exponent *θ*. In the optimization problem (4.8), we set *L*_1_ = 30 and *L*_2_ = 50. Table 1 reports the numerical estimates and standard deviations of *α, α* + *γ, τ* and *θ* averaged over a sample of up to 50 participants in the alert and fatigue states. For each participant, we use EEG measurements over 20 seconds at all 32 channels on the scalp. Fig. 7 plots the paths of the estimated values of *α, α*+*γ* and *τ* over these participants.

**Figure 4:**
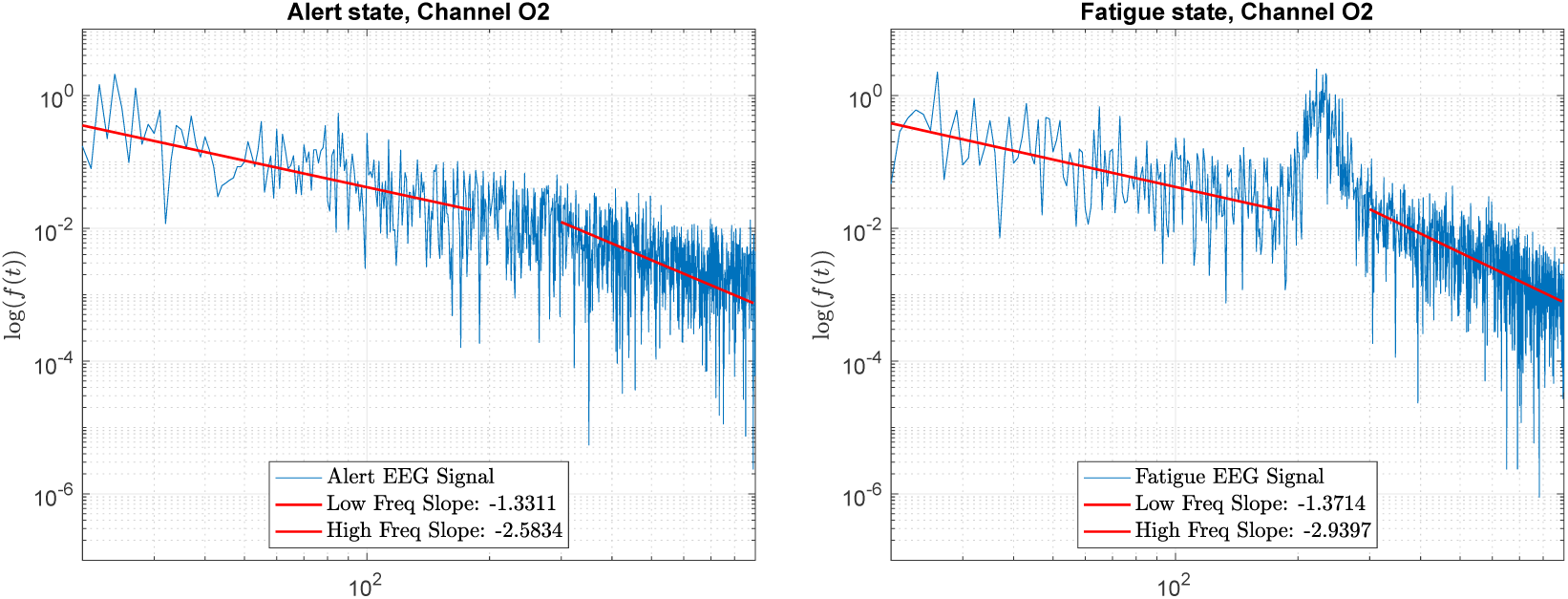
Log-periodogram of the EEG time series at channel O2 shown in Fig. 3

**Figure 5:**
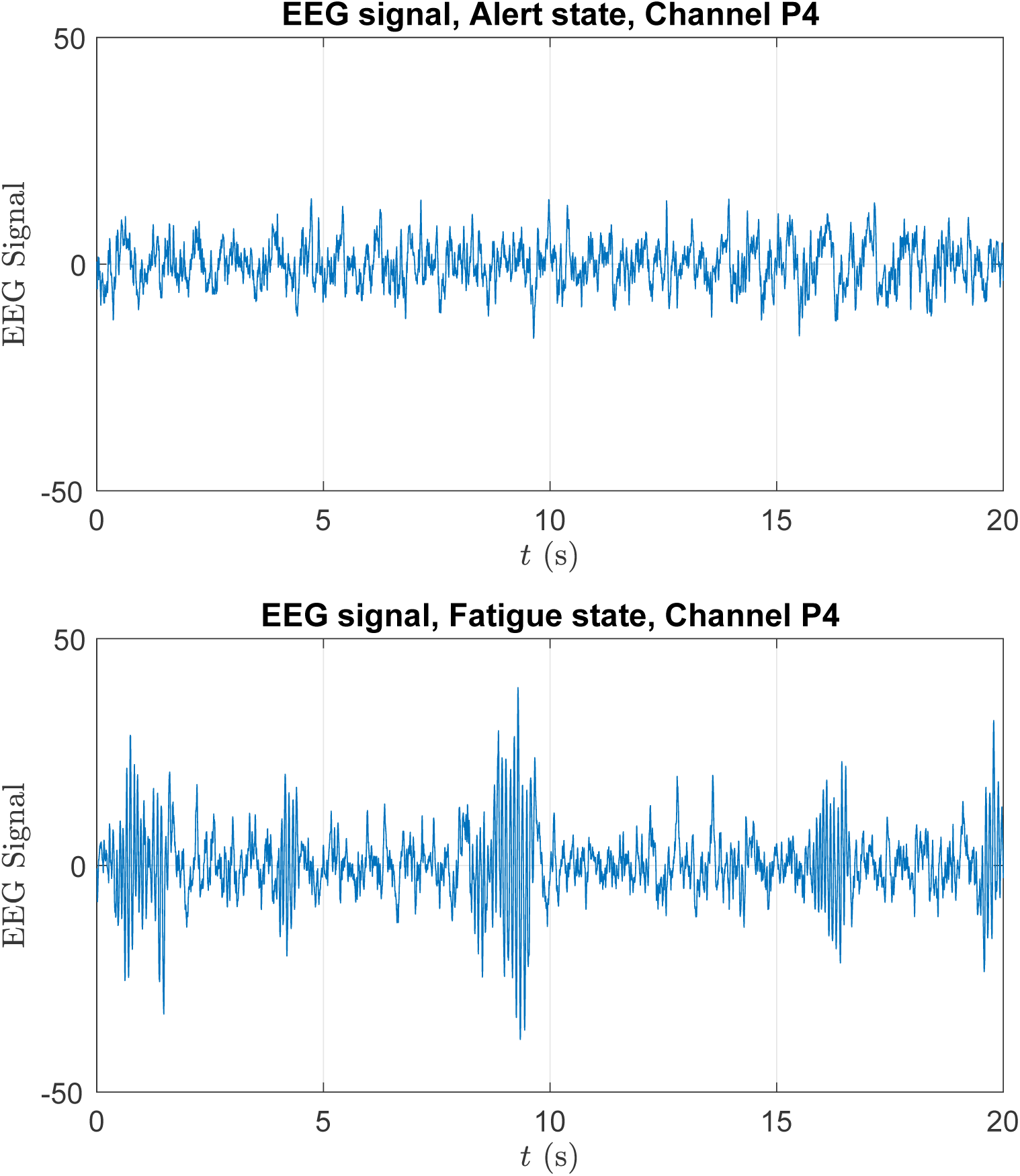
EEG time series at channel P4

**Figure 6:**
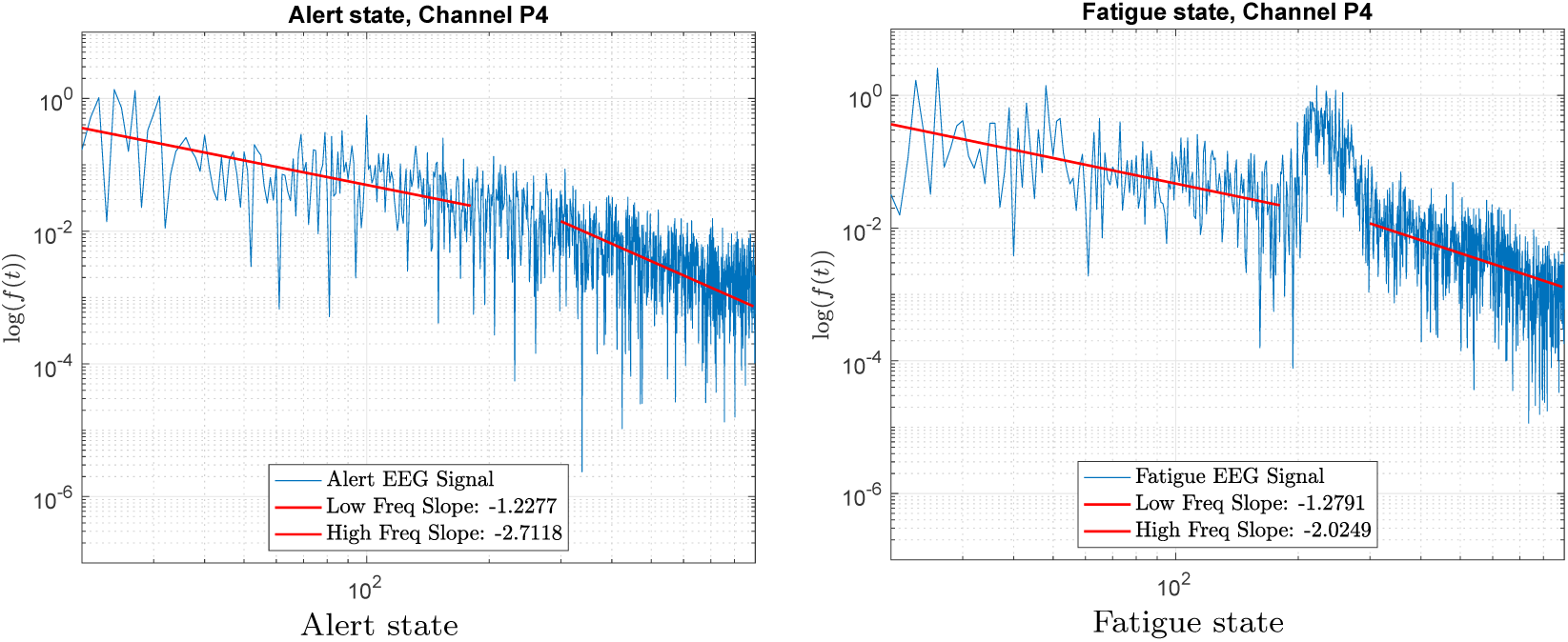
Log-periodogram of the EEG time series at channel P4 shown in Fig. 5

**Figure 7:**
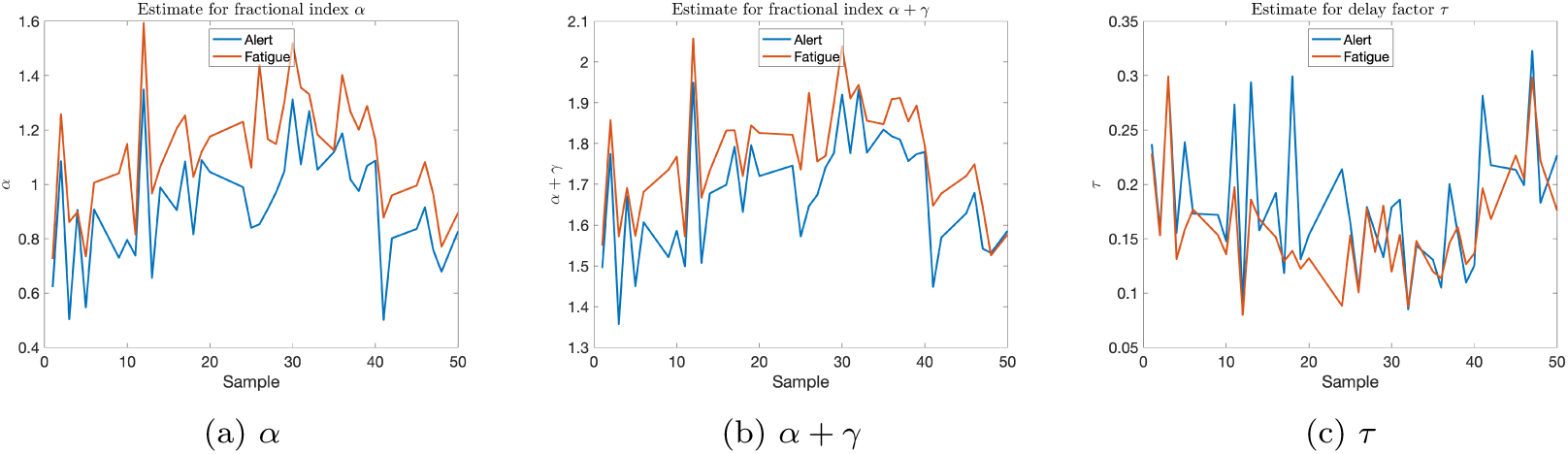
Paths of fractional diffusion exponents *α, α* + *γ* and delay parameter *τ* over a sample of up to 50 participants

First of all, the averaged value *α* = 1.116 in Table 1 for the fatigue state yields *β* = 0.21, hence *κ* = 0.79 in the spectral density (5.2). This result indicates that global EEG exhibits long memory, hence global power-law scaling in the fatigue state, as predicted by the covariance function (5.1) and the spectral density (5.2). The path of *α* in Fig. 7 also shows that *α* is consistently larger than 1 over the sample of participants, hence exhibiting power-law scaling in the fatigue state. The averaged value of *α* smaller but close to 1 for the alert state in Table 1 does not lend support for the assertion of long memory in the alert state. This would indicate weakly dependent non-stationarity rather than strongly dependent stationarity in the solution for the alert state. This agrees with the assertion of higher non-Gaussianity in the alert state, which we discuss next.

The estimates of *α* and *α* + *γ* are obtained for *t* sufficiently large. The value of *α* around 1 in Table 1 confirms that global EEG is *α*-stable in both alert and fatigue states. This degree of non-Gaussianity is distinct from the Gaussianity of standard diffusion when *α* = 2. It demonstrates the fractal effect due to a multifractal medium on the diffusion as anticipated in Eq. (2.11). The result agrees with the diffusion of EEG signals through a heterogeneous medium due to the flow of current in a conductive fluid and the high density of membranes as heralded in [7]. The lower value of *α* in Table 1 and consistently over the entire sample of participants in Fig. 7 for the alert state indicates that global EEG has larger jumps and more rugged paths in the alert state than in the fatigue state. The lower value of *α* + *γ*, in Table 1 and Fig. 7, corroborates the assertion that paths of global EEG are more multifractal in the alert state than in the fatigue state. All these interpretations are suggested by the analytical results of the fractional diffusion model of Subsection 2.3.

The mean value of the delay parameter *τ* in Table 1 is obtained under the assumption that the delay response is due to the initial condition, that is, when the alert or fatigue state starts. Hence *τ* is evaluated as *t →* 0 in the estimation scheme. The larger value of *τ* shown in Table 1 indicates that global EEG has longer delay, hence stronger memory, in the alert state than in the fatigue state. This result is also maintained consistently over the sample of participants in Fig. 7.

The above analysis confirms the occurrence of strong response of the system to the initial state within the context of non-Gaussian diffusion. The occurence is intuitively consistent with the behaviour of a driver in the alert or fatigue state. The results provide additional tools to construct global indicators to distinguish between these two states of brain activity complementing those afforded by time-series techniques.

## Acknowledgements

Hung T. Nguyen acknowledges support from the Australian Research Council under Discovery Project DP150102493. Yu Guang Wang acknowledges the support of funding from the European Research Council (ERC) under the European Union’s Horizon 2020 research and innovation programme (grant agreement n° 757983).

## Appendix A. Fundamental solution of delay-differential equations

In order to derive the solution of the system (1.1) and (1.2), we recall the following theory of fundamental solution of delay-differential equations in Banach spaces due to [30]. Let *E* be a Banach space. We denote by *B*(*E*) the Banach space of all bounded linear operators from *E* into itself. We consider the following differential system with *m* delay terms:

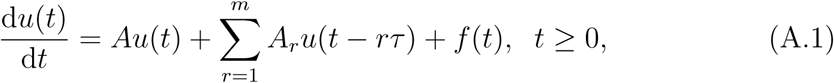

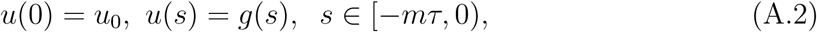

where *τ >* 0 is a constant, *u*(*t*), *f* (*t*), *g*(*t*) *∈ E*. The operators *A* and *A*_*r*_, *r* = 1, *…, m*, possibly unbounded, are assumed to satisfy the following assumptions:

**H**_0_. The operator *A* generates a strongly continuous semigroup {*T* (*t*), *t ≥* 0} on *E*;

**H**_1_. The *A*_*r*_, *r* = 1, *…, m*, are closed linear operators with dense domains *D*(*A*_*r*_) in *E*;

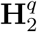. For each *r*, there exists a function *M*_*r*_(*·*) *∈ L*_*q*_([0, *τ*]) such that *∥T* (*t*)*A*_*r*_*u∥ ≤ M*_*r*_(*t*)*∥u∥* for almost all *t ∈* [0, *τ*] and all *u ∈ D*(*A*_*r*_).

Let 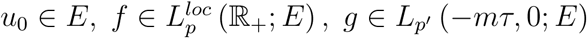, be given, and let Assumption 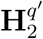 with 1*/p′*+ 1*/q′*= 1 be satisfied, where *p, p′∈* [1, *∞*]. Then, the function

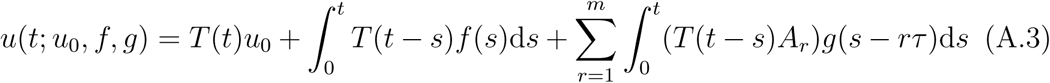

is well defined and is strongly continuous on [0, *τ*]. This function is defined to be the mild solution of the system (A.1) and (A.2), see [30].

Let now *f* = 0, *g* = 0 and let Assumptions **H**_0_, **H**_1_ and 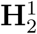 be satisfied. Then, as given by (A.3), the mild solution *u*(*t*; *u*_0_) = *u*(*t*; *u*_0_, 0, 0) can be constructed for any *u*_0_ *∈ E*. The mapping *𝒢* : ℝ_+_ *× E → E* defined by *G*(*t, u*_0_) = *u*(*t*; *u*_0_) generates a one-parameter family of bounded operators {*G*(*t*), *t ≥* 0}, where *G*(*t*) is defined by *G*(*t*)*u* = *G*(*t, u*) for *u ∈ E* and satisfies

i. *G*(*t*) = *T* (*t*) for all *t ∈* [0, *τ*] and *G*(*t*) *∈ B*(*E*) for all *t ≥* 0;
ii. for each *u*_0_ *∈ E, G*(*t*)*u*_0_ is continuous on ℝ_+_.

By [30], we can define the function *G*(*t*) as the fundamental solution of Eq. (A.1). From the given semigroup {*T* (*t*), *t ≥* 0} we define the operators *T*_1_(*t*), *…, T*_*k*_(*t*), *t ≥* 0, inductively by

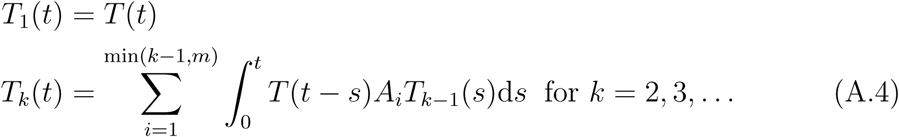

From (A.4), an explicit form of *T*_*k*_(*t*) can be derived. We first define the index sets Λ(*j, k*) for all *j, k* = 1, 2, *…* by

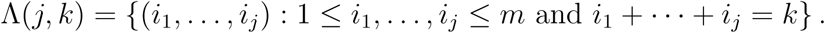

Note that Λ(*j, k*) = ∅ for *j > k*. The following integral expression of *T*_*k*_(*t*) for *k ≥* 2 is then obtained:

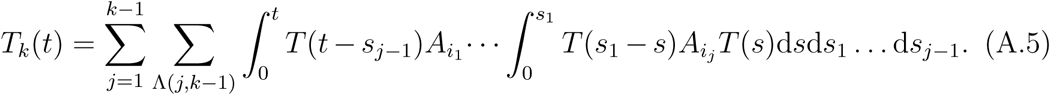

Summarising, we obtain the following result, which is given by [30, Theorem 3.1].

### Theorem Appendix A.1.

*Let Assumptions* **H**_0_, **H**_1_ *and* 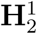 *be satisfied. Then the set of one-parameter family of strongly continuous operators* {*T*_*k*_(·) : *k* = 1, 2, *…*} *can be constructed as in* (A.5), *and the fundamental solution G*(*t*), *t ≥*0, *is given by*

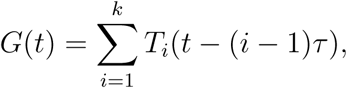

*where t ∈ [(k − 1) τ; kτ]*.

We introduce an additional condition:

**H**_3_. For each *r, A*_*r*_ commutes with *T* (*t*) for all *t ≥* 0, i.e., for any *u ∈ D*(*A*_*r*_) and *t ≥* 0, *T* (*t*)*u ∈ D*(*A*_*r*_) and *A*_*r*_*T* (*t*)*u* = *T* (*t*)*A*_*r*_*u*.

Then, by **H**_3_ and (A.5), *T*_*k*_(*t*) is given formally by

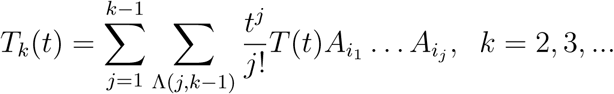

Consequently, *G*(*t*), *t ∈* [(*k* − 1) *τ, kτ*], is represented formally as

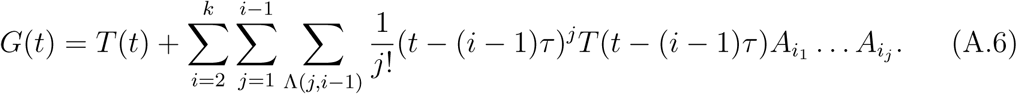

The mild solution (A.3) can then be given in terms of *G*(*t*) as

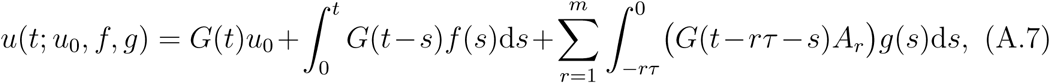

where *G*(*s*)*A*_*r*_ = *O*, the null operator on *E*, if *s <* 0. This is given by [30, Theorem 4.2].

